# Association between high-frequency hearing sensitivity and visual cross-modal plasticity in typical-hearing adults

**DOI:** 10.1101/2025.04.28.648945

**Authors:** Brandon T. Paul, Arunan Srikanthanathan, Maya Daien, Andrew Dimitrijevic

## Abstract

Visual cross-modal plasticity after deafness describes when auditory cortical regions increase responsiveness to visual stimulation. Such neuroplasticity demonstrates that sensory cortex adapts to profound peripheral impairment by upregulating healthy sensory function. However, whether cross-modal plasticity occurs for mild, partial hearing loss is less clear. Here we investigated cross-modal plasticity in “normal” hearing adults who had varying high-frequency hearing sensitivity, and if cross-modal plasticity was linked to adaptive visual behaviours. Twenty-five participants (aged 18 to 78) were tested in a modified Sternberg task with text-based visual words. Participants viewed a five-word sentence presented word-by-word during an encoding period and were asked to hold those words in short-term memory during a retention period. After, they reported if a target word probe was present in the encoding period. Multichannel EEG recorded visual cortical activity throughout. Results showed separate effects of high-frequency hearing sensitivity and age on visual cortical responses. At scalp sensors, visual N1 amplitudes were larger with worse hearing sensitivity, and P2 latencies were longer. In contrast, age was associated with shorter P2 latencies. Source analysis found that N1 amplitude increased with worse hearing sensitivity in right auditory cortex, but no effects were found in visual cortex, or for age. Hearing sensitivity and visual encoding activity did not correlate with behavioural performance or neural oscillations during the retention period. Results suggest for the first time that even a slight hearing decline in the high-frequency range is associated with higher visual activation of right auditory cortex, consistent with cross-modal plasticity.

## 1. Introduction

Losing a sense like hearing or vision can significantly impact everyday function, but the brain can compensate by relying on its remaining senses. Deaf individuals, for example, perform better than hearing controls on visual tasks that involve motion detection or visual localization (Codina et al., 2011; Shiell et al., 2014). Neuroscience research in animal models shows that these supranormal visual abilities are causally mediated by forms of neuroplasticity in sensory cortex (Lomber et al., 2010). “Cross-modal” plasticity occurs when brain regions deprived of their primary afferent input increase their sensitivity to external senses, evidenced in deafness when visual stimulation increasingly activates auditory cortical neurons (Finney et al., 2001; Lomber et al., 2010). Similarly, blind individuals show enhanced hearing function and visual cortex activation compared to sighed controls (Kujala et al., 1997; Rauschecker & Korte, 1993; Voss, 2016). It is argued that cross-modal plasticity is *compensatory* because it offsets deficits caused by the lack of hearing (Kral & Sharma, 2023; Lomber et al., 2010). Additionally, an “intra-modal” form of neuroplasticity occurs when sensory processing improves within its respective sensory cortex, such as when visual cortical activity increases in deafness (Bavelier et al., 2000).

A theoretically important question is whether mild, partial hearing loss leads to similar compensatory cross-modal or intra-modal plasticity. Most research on cross-modal plasticity investigates congenital deafness or profound hearing loss early in life where auditory cortex is entirely or mostly deprived of input. In contrast, partial hearing losses, such as noise-induced hearing loss and age-related hearing loss, are more common later in adulthood. In these cases, there is accumulation of damaging effects over the lifespan, caused by intense sound exposure, biological aging, ototoxic substances, and genetic factors (Huang & Tang, 2010). These partial hearing losses disproportionately affect higher frequencies of the hearing range, above 2 kHz, which increases in severity with age (Toppila et al., 2001). Because auditory cortical regions receive reduced, but nonetheless substantial input from the auditory periphery in partial hearing loss, neuroplasticity may unfold differently than conditions of total sensory deprivation.

Research in the past decade provided initial evidence for cross-modal plasticity in partial hearing loss. Most studies used visual stimuli such as speech, object movement, or shapes to evoke cortical potentials in the electroencephalogram (EEG). Visual-evoked potentials (VEPs) in those with mild-to-moderate partial hearing loss tend to have shorter latencies or larger amplitudes compared to those with typical hearing (Aguiar et al., 2025; Campbell & Sharma, 2014, 2020; Glick & Sharma, 2020; Liang et al., 2020). Larger VEP amplitudes with earlier latencies are consistent with cross-modal plasticity effects found in cochlear implant users with profound hearing loss (Doucet et al., 2006). Interestingly cross-modal plasticity in mild, partial hearing loss seems to be a reversible process because hearing aid use, which partially restores auditory input, was shown to restore visual cortical activation to levels found in typical hearing individuals (Glick & Sharma, 2020). Although cross-modal plasticity could explain these and other VEP effects found in adults with partial loss, intra-modal plasticity may also play a role. For instance, one functional magnetic resonance imaging (fMRI) study found increased functional connectivity within visual cortex in adults with age-related hearing loss (Ponticorvo et al., 2022), which could reflect intra-modal plasticity.

One unanswered question is if cross-modal plasticity is evident even when hearing loss is subtle. Studies across different forms of sensory loss, including EEG studies shown above, show that cross-modal plasticity does not require severe deafferentation and does not involve large-scale reorganization of sensory cortical regions (Makin & Krakauer, 2023). Instead, visual cross-modal plasticity in the auditory system likely results from synaptic scaling for “heteromodal” inputs in higher-order auditory regions (e.g., inputs from visual or somatosensory regions) that change dynamically based on the relative degree of auditory input (Kral & Sharma, 2023, Glick & Sharma, 2020). Past cross-modal research has focused on individuals with clinical evidence of mild to moderate hearing loss determined by measuring audiometric thresholds for pure tones across the hearing frequency range for speech (250 to 4000 Hz). However, studies reliably find associations between cross-modal plasticity and hearing loss in the high-frequency hearing range above 4 kHz (Aguiar et al., 2025; Glick & Sharma, 2020; Rosemann & Thiel, 2018). To test the assumption that cross-modal effects can be triggered by subtle losses, all participants in this study had conventionally “normal” hearing but exhibited variation in high-frequency hearing sensitivity, especially with increasing age. We carefully delineate aging effects from hearing effects for this reason.

A second unanswered question is whether cross-modal plasticity confers a behavioural advantage in individuals with partial hearing loss, as is found in deafness (Lomber et al., 2010). Based on past research in deafness, visual behavioural compensation should involve “supramodal” perceptions of objects that have both visual and auditory analogues; e.g., localizing an object in space, tracking its motion, or perceptions of speech and language (Kral & Sharma, 2023; Lomber et al., 2010). Compensatory behaviours will also likely involve increased recruitment of frontal cognitive systems, because cross-modal plasticity is shown to arise through top-down influences on higher-order auditory cortical regions (Berger et al., 2017; Yusuf et al., 2021), In agreement, EEG and fMRI studies have found frontal activity correlates with visual cross-modal responses in partial age-related hearing loss (Campbell & Sharma, 2020; Glick & Sharma, 2020; Rosemann & Thiel, 2018). However, to our knowledge, no studies directly show correlations between visual cross-modal plasticity in auditory cortex and changes to visual behaviours in partial hearing loss.

To this end, the present study examined visual cortical activity in individuals with varying hearing sensitivity while they were engaged in a cognitive short-term memory task. Stimuli were words in the format of visual text, and the task structure separated visual encoding processes from the retention of visual words in short-term memory, affording comparison of visual cortical potentials, behavioural outcomes, and cognitive processes. Visual text is an appropriate stimulus because it taps into supramodal lexical representations. In addition, those with worse hearing sensitivity may more often rely on closed captioning to supplement auditory speech understanding for video medias. We previously found evidence for enhanced visual responsiveness in cochlear implant users engaged in a similar Sternberg-style task with visual orthographic characters (Prince et al., 2021), but this design has not been evaluated in those with only mild changes to hearing sensitivity.

Specifically, we had four hypotheses:

H1: High-frequency hearing sensitivity will correlate with shorter VEP latencies and larger VEP amplitudes, consistent with cross-modal plasticity. In addition, source analysis will determine if differences localize to auditory or visual cortical regions.

H2: Cortical oscillations during the short-term memory retention will correlate with high-frequency hearing sensitivity.

H3: Visual cortical activity in those with worse hearing sensitivity will predict improved performance on the memory task.

H4: Participants’ numerical age will show distinct relationships to visual cortical activity that are separate from effects of hearing sensitivity, consistent with past research (Aguiar et al., 2025).

## 2. Methods and materials

### 2.1 Participants

Twenty-five participants were recruited from participant databases as Sunnybrook Health Sciences Centre. 18 to 78 (M = 41, SD = 19.7) and included 13 females and 12 males (self-reported). No participant indicated mental health or neurological conditions. Participants provided written informed consent, and the protocol of the study was approved by the Research Ethics Board at Sunnybrook Health Sciences Centre. The procedures were in accordance with the Declaration of Helsinki. Participants received monetary compensation of $25 CAD for completing the study, and if they traveled to the hospital campus in their personal vehicle, they received reimbursement for parking fees.

### 2.2 Hearing thresholds

We assessed hearing sensitivity using pure-tone audiometry, the gold standard test for hearing loss that obtains 50% detection thresholds for pure tones of varying frequency. Thresholds are measured in decibels hearing level (dB HL) and are higher when hearing sensitivity is worse. The experimenter used the Hughson-Westlake procedure to estimate thresholds for pure tones ranging from 250 Hz to 8 kHz in octave intervals. Measurements were made on a calibrated Grason-Stadler GSI 61 audiometer using TDH-39 supra-aural headphones within a sound-attenuated and electrically shielded sound booth. There is no international standard marking the degree of threshold shift at which hearing loss begins. One established convention takes an average of threshold estimates from 250 Hz to 4 kHz in octave steps. If this threshold average exceeds 25 dB HL in the better ear (better-ear pure-tone average, BEPTA), a person is considered to have evidence of hearing loss (Olusanya et al., 2019). The BEPTA averaged 7.84 dB HL in this sample (SD = 7.54, range –3 to 26 dB HL). Two individuals presented with a BEPTA of 26 dB HL, which slightly exceeds the range of typical hearing. Since this elevation was minor, we consider all participants to have hearing thresholds within a normal range.

One drawback to the BEPTA is that hearing thresholds are more likely to vary at higher frequencies above 2 kHz across the lifespan. Studies commonly estimate hearing sensitivity above the BEPTA range when demonstrating correlations to visual cortical activity using EEG (Aguiar et al., 2025; Glick & Sharma, 2020), fMRI (Rosemann et al., 2020), or for general health outcomes (Ramage-Morin et al., 2019). Following this convention, we take a cross-ear average of 4 and 8 kHz thresholds as a continuous measure of high-frequency hearing sensitivity.

### 2.3 Stimuli and task design

Visual stimuli were presented to participants in a modified verbal Sternberg task (Sternberg, 1966). In each trial, participants viewed a five-word sentence word-by-word, and after a brief interval, reported of a final target word was in the original sentence on that trial. Sentences used in the task were taken from the Matrix Sentence Test (Hagerman, 1982) shown in Figure 1A. The matrix sentence test is typically in an audio format but were converted to text characters for this study. Each sentence consists of 5 words in the same syntactic structure (subject, verb, number, adjective, direct object). For each sentence, words are randomly drawn in each position from 8 options, totaling 40 possible words. Words were presented as white text on a black background on a computer monitor spaced 1.5 m from the participant.

**Figure 1.**
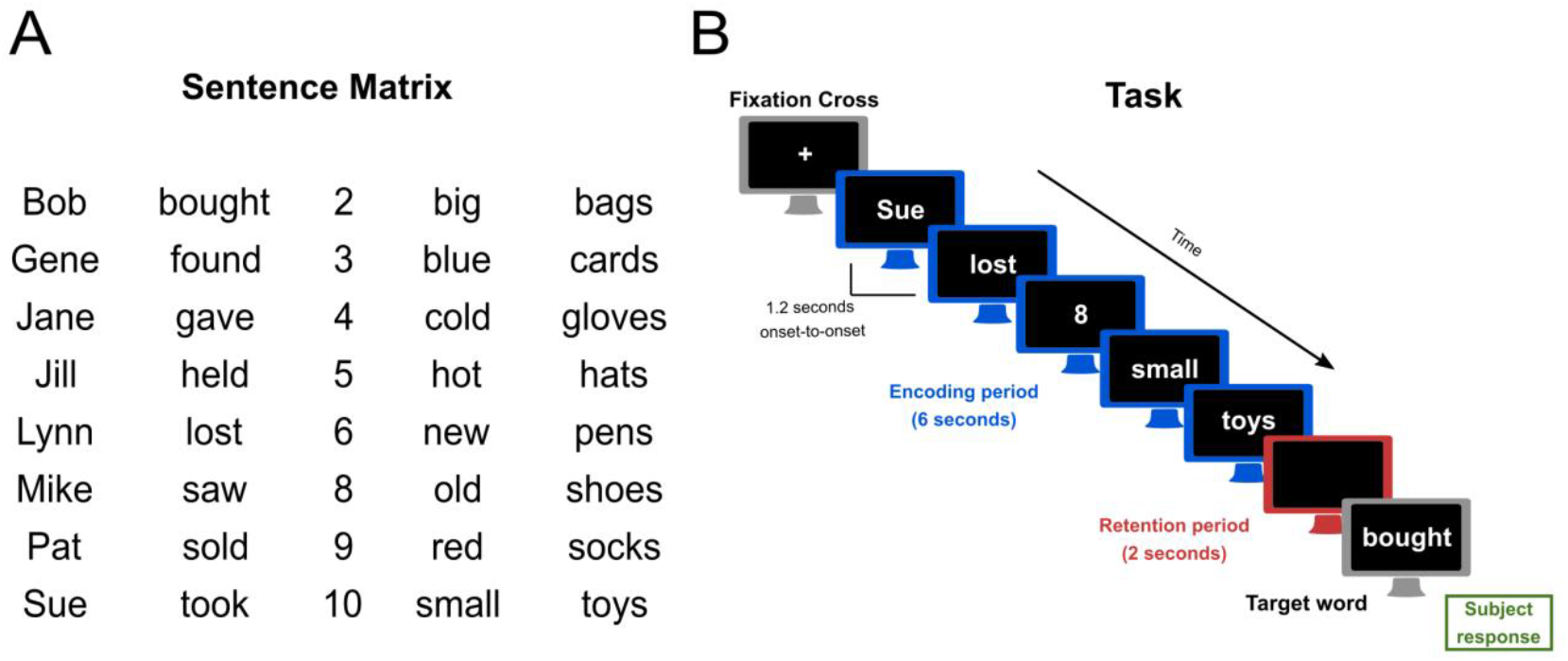
Stimuli and task. A) Sentence matrix showing the possible words that could appear during the trial. B) Task structure showing the presentation of words on a computer screen during an encoding period (blue), the retention of those words in short-term memory (red), followed by a target word. Participants reported by keypress whether target words occurred in the original sentence (green).

The structure for each trial is presented in Figure 1B, and consists of encoding, retention, and testing intervals. First, participants viewed a fixation cross in the center of the computer screen for 2 seconds. After, the encoding interval began after, and each word of the 5-word sentence was presented on the screen for 1.2 seconds and stayed on the screen until the onset of the next word. Words did not disappear in between word onsets. After termination of the final word, a black screen appeared for 2 seconds, which marked the retention interval where participants are presumed to maintain the sentence in short-term memory. After the retention interval, a target word appeared on the screen. The target word either appeared in the 5-word sentence of that trial or was randomly selected from the remaining words of the sentence matrix. The target word had a 50% probability of either outcome. Participants then reported if the target word was in the original sentence by pressing the left arrow key on a keyboard for “yes,” and the right arrow key for “no.” Reaction times were also recorded relative to the onset of the target word, but participants were not explicitly instructed to respond as quickly as possible. Participants received no feedback. Participants practiced this task for 10 trials before the study session began.

### 2.4 EEG Recording and Preprocessing

During the working memory task, the 64-channel EEG was sampled at 2000 Hz using a Neuroscan amplifier (Compumedics, Victoria, Australia) adapted for an actiChamp Brain Products cap (Brian Products GmbH, Inc., Munich, Germany). The channel layout used an equidistant montage that improves source localization, as reported in our previous studies on cross-modal plasticity in cochlear implant users (Paul et al., 2022, 2025; Prince et al., 2021). Channel Cz was the on-line reference, and the ground was 50% of the distance from Cz to the nasion. A Polhemus FastTrack digitizer (Colchester, USA) recorded the 3-dimensional sensor positions to a text file for later source analysis. Preprocessing was completed in Brain Vision Analyzer software v2.0 (Brain Products GmbH Inc., Munich, Germany). Data were first high pass filtered at 0.1 Hz to remove baseline drift, and downsampled to 250 Hz. Artifacts exceeding 500 mV were visually identified and marked for removal. Independent components analysis (ICA) decomposed continuous EEG to identify and remove eyeblinks and other myogenic artifacts. Continuous data were then exported to MATLAB.

### 2.5 Visual evoked potentials during encoding

We used the Fieldtrip Toolbox (Oostenveld et al., 2011) to analyze visual evoked potentials to visual word onsets in MATLAB. Continuous data were first imported from Brain Vision Analyzer. Bad channels were replaced with estimates based on neighbouring channels using spline interpolation. Trials with voltage fluctuations that were > 3 standard deviations of the mean were considered artifactual and were removed. After, data were epoched from 100 ms prior to word onset to 400 ms after word onset and collapsed across each of the word positions. Cleaned data were then referenced to the common average, and baseline corrected from –100 to 0 ms. Based on inspection of grand average evoked potentials, 4 channels in parietal and occipital regions were selected for VEP analysis (see Results for channel locations). Latencies for P1, N1, and P2 visual evoked potentials were selected as the time when voltages reached a maximum (negative maximum for N1). Amplitudes were computed as the average of a 16 ms window around the voltage maximum latency for each component.

### 2.6 Source modeling of visual evoked potentials

Standardized low-resolution electromagnetic tomography (sLORETA)(Palmero-Soler et al., 2007; Pascual-Marqui, 2002) was used to estimate sources of visual evoked potentials, via default settings in the Brainstorm toolbox in MATLAB (Tadel et al., 2011). Boundary element model (BEM) head models were constructed using the OpenMEEG plugin in Brainstorm (Gramfort et al., 2010; Kybic et al., 2005). The absolute values of the model activations were z-scored relative to the baseline. Visual and auditory cortical sources were extracted into regions of interest (ROI). from the Desikan–Killany atlas (Desikan et al., 2006). Following source modelling practices from Strophal and Debener (2017), Heschl’s gyrus was set as the auditory ROI. The pericalcarine cortex (primary visual) was the visual ROI. Analysis was performed on the ROIs’ time-waveform activation on a 16-ms time window centered on the N1 and P2 maxima.

### 2.7 Time-frequency analysis

We used time-frequency analysis to examine the power of neural oscillations during the trial period. Continuous EEG data were first referenced to the common average, epoched from −1 to 10.5 s relative to the onset of the first word and subjected to time-frequency analysis in Fieldtrip. Morlet wavelets with 5 wave cycles were used to estimate time-frequency representations in 1 Hz steps from 1 to 30 Hz. After, power values were expressed as a decibel change from baseline.

### 2.8 Statistical analysis

Statistical tests were conducted in R (R Core Team, 2021) and using built-in functions of the Fieldtrip toolbox in MATLAB. All tests were two-tailed, and the alpha criterion for Type I error was set at 0.05. It is important to note that age and high-frequency hearing thresholds were significantly correlated in this study (r = 0.63, p < 0.001) requiring caution when performing statistical models due to the chance of problematic collinearity between these variables that could bias slope coefficients. Importantly, variance inflation factors, which describe multicollinearity in linear models, were below 2.6 for both age and high-frequency hearing thresholds in the models described below, where values above 5 indicate potentially unacceptable multicollinearity. Therefore, it is unlikely that age and hearing thresholds are confounded or substantially bias the model estimates.

#### 2.8.1 Behavioural data

Logistic mixed effects regression was used to predict single-trial performance as a function of age and high-frequency hearing thresholds. Inverse gamma models were used to predict single-trial reaction times as a function of age and hearing thresholds. No evidence of model misspecification was found after inspecting residuals and QQ plots using the DHARMa package in R.

#### 2.8.2 Visual evoked potentials during the encoding period

Linear models assessed the effects of high-frequency hearing thresholds and age in years on P1, N1, and P2 amplitudes and latencies during the encoding period. No correlations were found between P1 responses and hearing sensitivity, so we compared N1 and P2 responses in a mixed effects model that set VEP component as a within-participants fixed effect. This model tests if hearing thresholds and aging have different effects for visual cortical processing stages (earlier = N1, later = P2). Before modeling, N1 amplitudes were multiplied by –1 to invert measurements to positive values for more direct comparison to P2 amplitudes. The model structure used was:

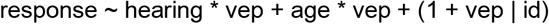

where *response* is either VEP amplitude in microvolts or latency in milliseconds, *hearing* is the 4–8 kHz hearing threshold in dB HL, *age* is participant age in years, *id* refers to participant ID code, and *vep* is a two-level, deviation-coded categorical variable representing the N1 or P2 component. Separate models were run for amplitude and latency. Model diagnostics revealed no aberrant indicators. Analysis of Variance (ANOVA) with Satterthwaite adjustment for degrees of freedom was used to test the significance level of model terms. Interactions were interpreted using simple slopes analysis.

#### 2.8.3 Visual evoked potential source analysis

Mixed models were also used to compare source activation magnitude and latency for visual and auditory ROIs. Separate models were run for N1 and P2 components. The model formula was:

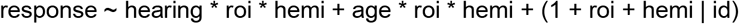

*Response* is either source latency or amplitude values, *hearing* and *age* terms were the same as the mixed model described for scalp VEPs, *roi* refers to the auditory or visual ROI, and *hemi* refers to the left or right hemisphere. The model predicts the neural response from three-way interactions between ROI, hemisphere, and hearing thresholds or age, while accounting for random effects including the random intercept for participant code (*id*), and random slopes for ROI and hemisphere. Separate models were run for amplitude and latency. Model diagnostics revealed no aberrant indicators. Analysis of Variance (ANOVA) with Satterthwaite adjustment for degrees of freedom was used to test the significance level of model terms. Interactions were interpreted using simple slopes analysis. Due to a technical problem with the EEG data file, one participant was removed from analysis, leaving 24 participants in the time-frequency analysis.

#### 2.8.4 Time frequency responses during retention period

We used Pearson correlations to test for associations between age and high-frequency hearing thresholds with neural oscillation power during the retention interval that elapsed 6–8 seconds after initial trial onset. Correlation coefficients were collected for all time points, sensors, and frequencies (1–30 Hz in 1 Hz steps) Test outcomes were subjected to cluster-based permutation tests using the *Fieldtrip* toolbox to sidestep the multiple comparisons problem (Maris & Oostenveld, 2007). During this procedure, correlation coefficients whose values falling under the alpha criterion of 0.05 were first summed into clusters based on spatial, temporal, or spectral adjacency. These summed values were then compared to a distribution of test values randomly generated from the dataset using Monte Carlo approximation (5000 iterations). If the proportion of random partitions that were larger than the observed test statistics was greater than 0.05 (two-tailed), the result of the cluster test was considered statistically significant. Separate sets of analysis were completed numerical age and hearing sensitivity level. Due to a technical problem with the EEG data file, one participant was removed from analysis, leaving 24 participants in the time-frequency analysis.

## 3. Results

### 3.1 Hearing thresholds, age, and task performance

High-frequency hearing sensitivity ranged from –1.25 to 55 dB HL (M = 17.90, SD = 16.29) and was correlated with age (r = 0.63, p < 0.001). Performance on the working memory task was high but not at ceiling (M = 91%, SD = 13%), and the average reaction time was 1120 ms (SD = 238 ms). A logistic mixed effects model on single trial performance did not show any relationships to high-frequency hearing thresholds (p = 0.45) or age (p = 0.47).

### 3.2 Visual evoked potentials during encoding

Visual evoked potentials across the whole trial are plotted in the top row of Figure 2A, and the grand average VEP collapsed across each word presentation is shown in Figure 2B. Clear P1, N1 and P2 components were present for each participant. Topographical maps in Figure 2C show that the P1 and N1 response reach a maximum in occipital sensors, while the voltage distribution was slightly anterior for the P2 response. P1 latencies and amplitudes did not correlate with age or hearing loss (p’s > 0.08). A mixed model comparing N1 and P2 amplitudes as a function of age and high-frequency hearing thresholds found no main effects for age, hearing thresholds, or VEP component (p’s > 0.28). There was no interaction between age and VEP component (p = 0.16). However, the interaction between VEP component and high-frequency hearing thresholds was significant (β = –0.11, SE = 0.05, F(1, 22) = 4.51, p = 0.045). Simple slopes analysis plotted in Figure 3A found that N1 amplitudes increased with higher high-frequency thresholds (i.e., worse hearing sensitivity) (β = 0.06, SE = 0.02, p = 0.029) but no relationship was found for P2 amplitudes (β = –0.05, p = 0.11).

**Figure 2.**
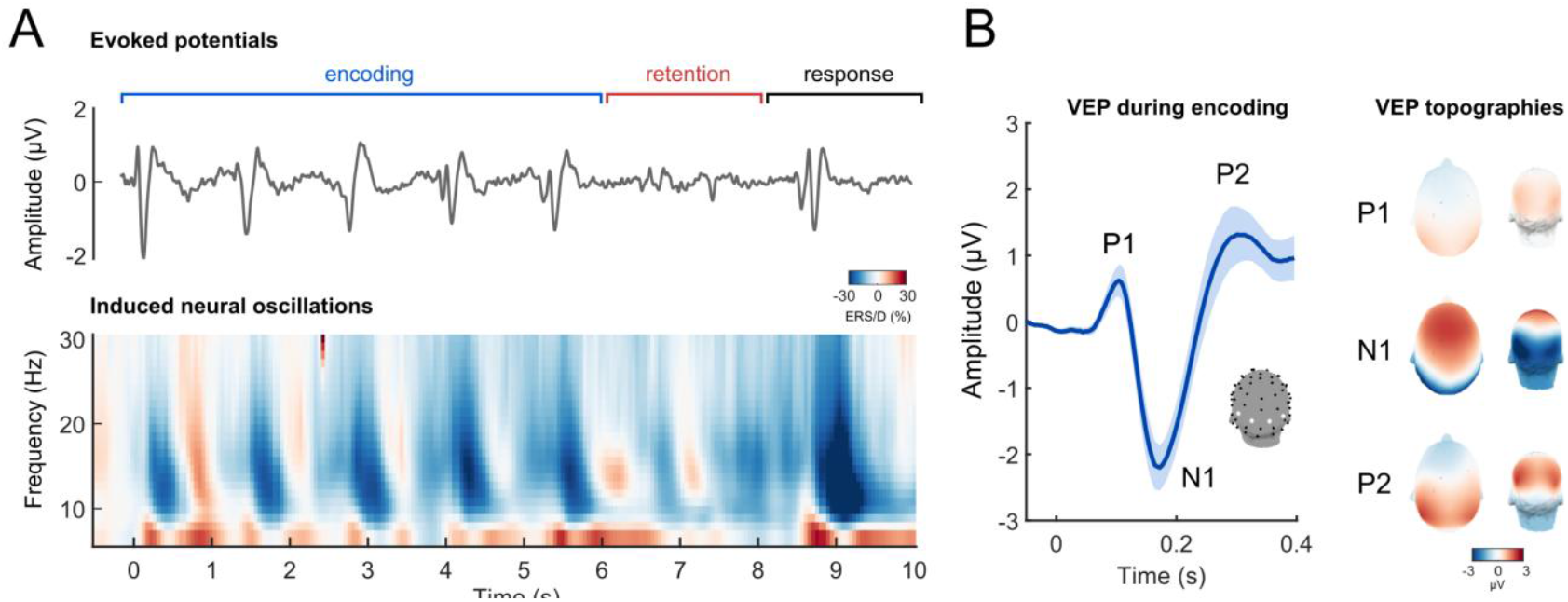
Visual cortical activity during the short-term memory task. A) The top row shows the time course of visual evoked potentials from occipital sensors across the trial. Encoding, retention, and response periods are outlined by brackets above the voltage waveform. The bottom row shows a spectrogram for the time course of induced neural oscillations, expressed as a normalized difference from baseline (event-related synchronization, ERS in red; event-related desynchronization, ERD, in blue). B) The left panel shows an average visual evoked potential over all characters presented in the encoding period. Clear P1, N1, and P2 responses are shown in occipital sensors. The right panel shows 3D topographic maps for P1, N1, and P2 potentials from occipital and dorsal perspectives.

**Figure 3.**
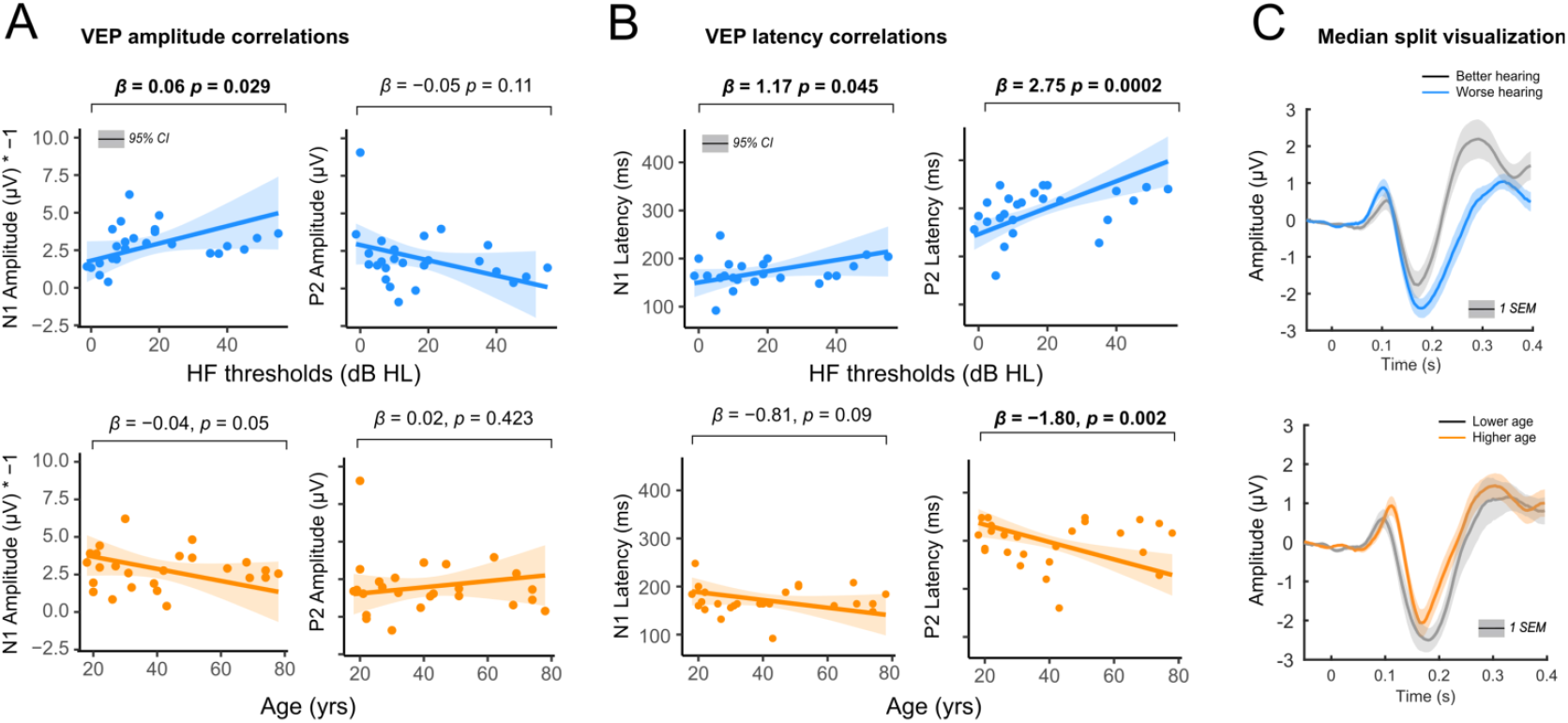
Correlations between high-frequency hearing thresholds, age, and visual evoked potentials during encoding. A) Scatterplots showing associations between N1 and P2 amplitude for high-frequency hearing (blue) and age (orange). Slope coefficients (beta values) and their p values are shown as a bracket above each plot. The shaded region represents the 95% confidence interval (CI). Significant associations are displayed in bold typeface. B) Same as panel A, but for VEP latencies. C) VEPs depicted for participants split at the median for high-frequency hearing thresholds on the top row, with worse high-frequency sensitivity shown as blue, and better sensitivity shown as grey. On the bottom row, VEPs separated by median-split age. Higher age is shown as orange, and lower age is shown as grey. Shaded regions represent 1 standard error of the mean (SEM).

For latency, there was a main effect of VEP component (β = 136.67, SE = 15.12, F(1, 22) = 81.63, p < 0.001), which makes sense given that P2 responses are ∼ 100–140 ms after N1 responses. A main effect of age (β = –1.31, SE = 0.49, F(1, 22) = 7.13, p = 0.014) suggested that P2 latencies were ∼13 ms shorter per each decade increase in age, and a main effect of hearing thresholds (β = 1.96, SE = 0.60, F(1, 22) = 10.80, p = 0.003) indicated an almost 20 ms increase in latency with 10 dB increase hearing thresholds. There was a significant interaction between VEP component and hearing thresholds (β = 1.58, SE = 0.61, F(1, 22) = 6.79, p = 0.016). Simple slopes analysis found that higher hearing thresholds (worse sensitivity) were associated with longer N1 latency (β = 1.17, SE = 0.55, p = 0.045) and P2 latency (β = 2.75, SE = 0.67, p < 0.001). The interaction between VEP component and age was not significant (p = 0.06). Simple slopes to understand separate VEP effects found no correlation between age and N1 latency (β = –0.81, p = 0.09) but shorter latencies were found with increasing age (β = –1.80, SE = 0.55, p = 0.002).

In a summary illustration, VEPs are shown dividing age and high-frequency hearing sensitivity as a median split in Figure 3C. Consistent with correlation analysis, N1 amplitude appears larger with worse hearing sensitivity, and the P2 had longer latencies. With higher age, N1 responses appear smaller with shorter latencies. These plots support the conclusion for separable effects of age and hearing sensitivity on visual evoked potentials. Note that inferential analysis was done modeling age and hearing sensitivity as continuous variables, not as dichotomous groups split by medians; this visualization is intended to show age and hearing sensitivity effects on VEP morphology overall.

### 3.3 Source analysis of visual evoked potentials

Source analysis was used to determine if visual cortical differences in those with worse hearing sensitivity resulted from intra-modal or cross-modal plasticity. Source activation for left and right auditory and visual cortex is shown in Figure 4. Mixed model analysis for the N1 source found no main effects. However, there was a three-way interaction between hearing thresholds, hemisphere, and sensory cortex (β = 1.31, SE = 0.61, F(1, 39.58) = 4.66, p = 0.037). To interpret this interaction, two further post-hoc mixed models were built for the left and right hemispheres and examined two-way interactions between hearing thresholds and sensory cortex. No significant main effects or interactions were found for the left hemisphere (p’s > 0.52). For the right hemisphere, the model returned only a two-way interaction between sensory cortex and hearing thresholds (β = 1.34, SE = 0.64, F(1, 21) = 4.44, p = 0.047). Simple slopes analysis found increased N1 activation magnitude with higher hearing thresholds in the auditory cortex (β = 1.49, SE = 0.56, p = 0.015) but not visual cortex (β = 0.14, SE = 0.80, p = 0.86). No main effects or interactions were found for N1 source latency (p’s > 0.14), P2 source magnitude (p’s > 0.26), or P2 source latencies (p’s > 0.27). In sum, results showed larger N1 cortical activation with worse hearing sensitivity that was specific to the right auditory cortex. No effects or interactions were found for age.

**Figure 4.**
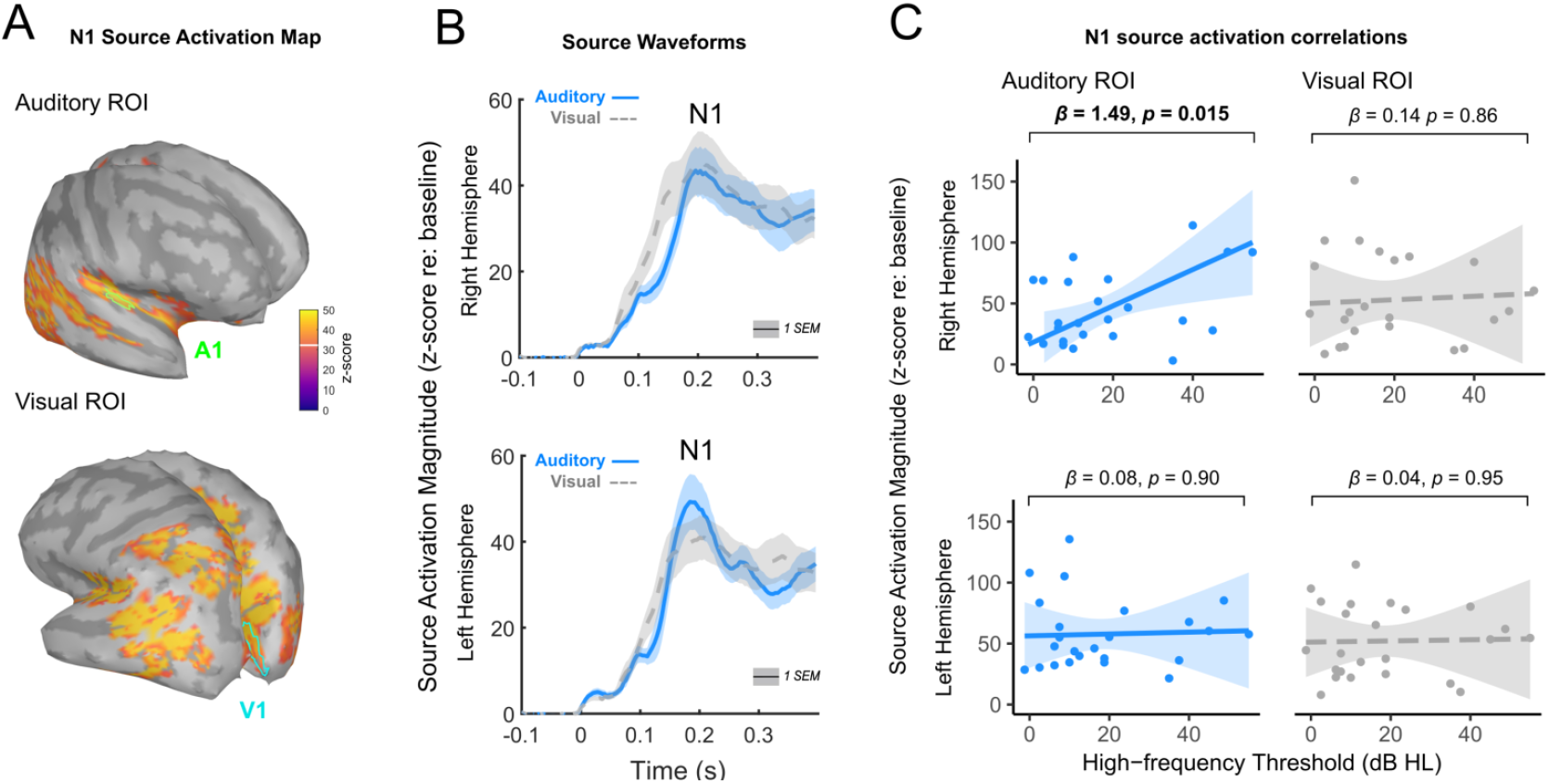
N1 Source activation and correlations between high-frequency hearing thresholds during encoding. A). Brain maps showing cortical activation during the N1 peak. The top row shows the location of the auditory ROI in green, and the bottom row shows the visual ROI in blue. B) Time series of source activations for the right hemisphere (top row) and left hemisphere (bottom row). Solid lines in blue represent the auditory source, and dashed lines in grey represent the visual source. The N1 peak is shown at ∼ 0.2 s. The shaded region represents 1 standard error of the mean (SEM). C) Scatterplots showing associations between high-frequency thresholds and N1 source activation for auditory ROIs (blue) and visual ROIs (grey). N1 source activation for the right hemisphere is shown in the top row, and the bottom row shows the left hemisphere Slope coefficients (beta values) and their p values are shown as a bracket above each plot. The shaded region represents the 95% confidence interval (CI). Significant associations are displayed in bold typeface.

### 3.4 Time-frequency analysis during retention

Induced power across the entire trial period is shown in in the bottom row of Figure 2A. The retention period was marked by two bursts of high alpha activity (10–15 Hz) in parietal sensors. However, cluster-based permutation tests did not show any significant associations between neural activity during the retention period with age (p’s > 0.3) or hearing loss (p’s > 0.65). N1 and P2 amplitudes and latencies during the encoding period also did not correlate with neural oscillations during the retention period (p’s > 0.16).

### 3.5 Brain and behavior correlations

Logistic mixed effects models did not find that N1 or P2 amplitudes and latencies correlated with performance on the memory task, nor did source activations (p’s > 0.15). Similarly, reaction times did not correlate with VEPs or source activation (p’s > 0.14).

## 4. Discussion

### 4.1 Summary

We tested for evidence of visual cortical neuroplasticity in individuals comparatively poorer hearing sensitivity using EEG, and to determine if this neuroplasticity was compensatory, we compared visual cortical responses to memory-related neural oscillations and short-term memory behaviour. Consistent with Hypothesis 1, at the scalp level, we found that cortical responses to the onset visual words showed larger N1 amplitudes and delayed P2 latencies with lower hearing sensitivity. These effects resemble evidence for visual neuroplasticity in mild-to-moderate hearing loss. At the source level, worse hearing sensitivity correlated with increased N1 activation in right auditory cortex, but no similar effects were found in visual cortex or for left auditory cortex. Consistent with Hypothesis 4, numerical age did not explain these relationships, and for the P2 response, analysis showed an opposite effect where latencies were shorter for increasing age. Inconsistent with Hypotheses 2 and 3, neither hearing sensitivity nor age correlated with memory accuracy, reaction time, or neural activity during the short-term memory task. Therefore, we found no evidence that visual neuroplasticity reflected a compensatory process, at least for the current task conditions and dependent measures.

### 4.2 Hearing loss, age, and visual-evoked cortical responses

All participants in this study qualified as having typical or near-typical hearing based on conventional “gold standard” criteria, defined as having audiometric hearing thresholds below 25 dB HL in the better ear averaged across the frequency range of 500 to 4000 Hz. However, hearing thresholds are more likely to vary with age at higher frequencies, such as at 4 and 8 kHz (Huang & Tang, 2010). This part of the hearing range and above is believed to be predictive of everyday listening ability such as of speech-in-noise perception, because high frequency information aids in extracting speech signals from a noisy environment (Zadeh et al., 2019). In addition, elevated high-frequency thresholds could indicate more extensive damage to cochlear synapses at lower frequencies that go undetected by threshold tests like the audiogram because they affect nerve fibers that code for suprathreshold sounds (Sergeyenko et al., 2013). Participants in this study with lower hearing sensitivity potentially experienced functional hearing difficulties to a degree that drove upregulation of visual processing, reflected in changes to N1 and P2 responses. Increased N1 amplitudes in those with worse hearing agree with past research that used a visual motion stimulus to show elevated N1 amplitudes in those with early-stage mild-to-moderate hearing loss. (Campbell & Sharma, 2014). Larger N1 responses may indicate an increase in the number of visual responding neurons, as has been found in multisensory areas in animal models of hearing loss (Schormans et al., 2017).

We also compared high-frequency hearing sensitivity to auditory and visual source activations during the N1 response. We found that hearing loss was correlated with a selective increase in right auditory cortical activation indicating cross-modal plasticity, but no differences were found in visual cortical regions or in left hemisphere that would indicate intra-modal plasticity. Right-lateralized cross-modal activation agrees with past studies in both profoundly deaf participants (Doucet et al., 2006) and in mild partial hearing loss (Glick & Sharma, 2020). Although Stropahl & Debener (2017) reported descriptively higher visual speech-evoked auditory cortical activation in participants with moderate haring loss that did not reach statistical significance, our findings are consistent with the direction of these effects.

For latency, we found that N1 responses were longer, not shorter with worse hearing sensitivity. This opposed initial predictions of visual plasticity following hearing loss and instead was consistent with known aging effects on VEPs. However, because P2 responses showed a similar effect of longer latencies, and this effect diverged from age which showed earlier P2 latencies, the findings suggest statistically separable influences of age and hearing ability. The visual P2 response is a late cortical potential measured at scalp sensors is a composite of neural generators in occipital, temporal and frontal areas (Freunberger et al., 2007), and is shown to be highly neuroplastic depending on a person’s sensory experience (Ahmadi et al., 2018). Visual P2 responses have consistently shown correlations to hearing function in with different stimulus types, suggesting it could be a useful neural marker for visual plasticity following hearing loss (Aguiar et al., 2025). However, the direction and spatial location of P2 effects have not been consistent in the literature.

Differential contributions from cortical generators that sum at the scalp level, which further depend on the type of visual events they process (e.g., speech, text, motion), could explain discrepant results between studies. Aguiar et al. (2025) recently showed diverging aging and hearing effects on the P2 latency in response to visual speech stimuli: P2 latencies were shorter with worse hearing and longer with higher age. Other studies using a visual motion stimulus have found that hearing loss is associated with shorter P2 latencies in right temporal regions (Glick & Sharma, 2020), larger P2 amplitudes in occipital regions (Campbell & Sharma, 2014), or smaller P2 responses in frontal (Campbell & Sharma, 2020) or occipital regions (Aguiar et al., 2025). Smaller P2 responses have also been found in response to visual shapes (Loughrey et al., 2023) in individuals with moderate compared to mild hearing loss. These studies suggest that the effect of hearing ability on visual evoked potentials may be highly stimulus-specific and depend on relative contributions from underlying generators. We did not examine effects of multiple generators on P2 activation in source analysis, which could explain why no correlation between source P2 activation and hearing status emerged. Future research should resolve these discrepancies by modeling these separate generators as a function of stimulus type given the P2’s promise as a visuoplastic neural marker (Aguiar et al., 2025),.

We did not find correlations between high-frequency hearing sensitivity and visual P1 responses. Larger or earlier P1 responses have been found in other studies using a visual motion stimulus (Campbell & Sharma, 2014, 2020; Glick & Sharma, 2020) or abstract visual shapes (Loughrey et al., 2023). P1 effects resulting from hearing loss notably contrast with known effects of age on visual cortical potentials. Age consistently predicts longer P1 latencies, (Brown et al., 2019; Price et al., 2017; Sokol et al., 1981; Tobimatsu et al., 1993) suggesting age-related decline in central sensory processing. Therefore, like the P2 response, the P1 response may change along independent trajectories for age and sensory loss. However, to our knowledge, age and hearing loss have not shown separate P1 effects in the same participants, and so further research is required to understand how these factors shape early visual cortical activity.

### 4.3 Associations between visual function, cognition, and hearing ability

Cross-modal plasticity is presumed to have a compensatory effect where vision helps to offset deficits caused by worse hearing function. A seminal study in by Lomber and colleagues (Lomber et al., 2010) found that visual motion detection and visual localization were enhanced in deaf compared to hearing cats. When higher-order areas of the auditory cortex were temporarily deactivated using cooling loops, these visual perceptual enhancements were abolished. Therefore, Lomber et al. convincingly demonstrated that cross-modal plasticity causally mediates improvements in visual function under total sensory loss. A similar link between visual cross- or intra-modal plasticity and improved visual-only behavioural outcomes in adults with partial hearing loss have not been reported to the authors’ knowledge. Here, our results do not support the explanation that hearing-loss related changes to cortical activity evoked by visual words yields a short-term memory benefit. We believed this format was appropriate to test for visual processing differences because visual text aids such as closed captioning can improve speech comprehension when auditory processing is challenged by hearing loss or environmental factors. A limitation to our study was that task performance was high, but not at ceiling. Compensatory effects may only appear with greater task difficulty. Other studies (Loughrey et al., 2023; Rosemann & Thiel, 2020; Zhu et al., 2024) also attempted to link visual cortical activity to cognitive behavioural differences in participants with and without hearing loss, but no report found statistically significant results.

Beyond visual processing, compensatory benefits of cross-modal plasticity may be more apparent when audio and visual information is integrated. Past research suggests that in people with mild to moderate hearing loss, perception is weighted toward vision rather than audition in multisensory tasks. Adults with age-related hearing loss, for example, are affected more by visual distractors in an auditory task compared to auditory distractors in a visual task (Puschmann et al., 2014). Participants with age-related hearing loss are also more susceptible to the McGurk effect, where conflicting auditory and visual speech phonemes create illusory phonemic fusions between the two modalities (Rosemann & Thiel, 2018; Stropahl & Debener, 2017). In addition, brain imaging research shows that hearing loss is associated with increased recruitment of frontal brain areas during audiovisual speech perception (Rosemann & Thiel, 2018), and stronger functional connectivity between auditory and visual cortical regions (Puschmann & Thiel, 2017; Rosemann et al., 2020). Therefore, it is possible that partial hearing loss increases demand on the visual system during audiovisual perception and frontal cortical regions may coordinate an increase in connectivity within and between sensory areas, consistent with predictions of cross- and intra-modal plasticity. It is unclear if these neural changes have consequences for visual-only perception.

### 4.4 Limitations and conclusions

This study has limitations. First, EEG has limited spatial resolution for resolving cortical sources, so interpretations based on source analysis should be taken with caution. Second, participants were not screened for cognitive impairment before the study, but they also did not disclose cognitive impairments in subjective self-report on demographic forms. Third, we did not measure visual evoked potentials under task-free conditions which would better allow a comparison between cortical activity during and absent of cognitive demands. Future studies on visual compensatory plasticity should consider comparing cortical activity inside and outside of task constraints. Fourth, high-frequency hearing ability is correlated to speech-in-noise perception, and speech-in-noise perception correlates with visual evoked potentials (Campbell & Sharma, 2014). We did not measure speech ability in our participants and cannot confirm these relationships.

To conclude, we report first evidence that cortical potentials evoked by visual words were associated with participants’ high-frequency hearing thresholds, even when their hearing was at typical levels. N1 responses were larger with worse high-frequency hearing sensitivity, and P2 latencies were longer. Source analysis of the N1 showed higher activation of right auditory cortex in those with worse hearing sensitivity. These effects were statistically separate from participants’ numerical age, and provided evidence that even mild hearing losses may shift the balance of visual and auditory processing across sensory cortex. However, visual cortical activity and hearing sensitivity did not correlate with behaviour or neural activity during short-term memory maintenance. Further research is required to understand if visual plasticity in hearing loss has an adaptive benefit for visual perception.

